# Cuticle supplementation and nitrogen recycling by a dual bacterial symbiosis in a family of xylophagous beetles (Coleoptera: Bostrichidae)

**DOI:** 10.1101/2022.12.09.519726

**Authors:** Julian Simon Thilo Kiefer, Eugen Bauer, Genta Okude, Takema Fukatsu, Martin Kaltenpoth, Tobias Engl

## Abstract

Many insects engage in stable nutritional symbioses with bacteria that supplement limiting essential nutrients to their host. While several plant sap-feeding Hemipteran lineages are known to be simultaneously associated with two or more endosymbionts with complementary biosynthetic pathways to synthesize amino acids or vitamins, such co-obligate symbioses have not been functionally characterized in other insect orders. Here, we report on the characterization of a dual co-obligate, bacteriome-localized symbiosis in a family of xylophagous beetles using comparative genomics, fluorescence microscopy, and phylogenetic analyses. Across the beetle family Bostrichidae, all investigated species harbored the Bacteroidota symbiont *Shikimatogenerans bostrichidophilus* that encodes the shikimate pathway to produce tyrosine precursors in its severely reduced genome, likely supplementing the beetles’ cuticle biosynthesis, sclerotisation, and melanisation. One clade of Bostrichid beetles additionally housed the co-obligate symbiont *Bostrichicola ureolyticus* that is inferred to complement the function of *Shikimatogenerans* by recycling urea and provisioning the essential amino acid lysine, thereby providing additional benefits on nitrogen-poor diets. Both symbionts represent ancient associations within the Bostrichidae that have subsequently experienced genome erosion and co-speciation with their hosts. While *Bostrichicola* was repeatedly lost, *Shikimatogenerans* has been retained throughout the family and exhibits a perfect pattern of co-speciation. Our results reveal that co-obligate symbioses with complementary metabolic capabilities occur beyond the well-known sap-feeding Hemiptera and highlight the importance of symbiont-mediated cuticle supplementation and nitrogen recycling for herbivorous beetles.

**Significance statement:** Nutritional symbioses evolved frequently in insects and contribute diverse metabolites to their hosts’ physiology. Associations with dual symbionts providing complementary nutrients evolved in multiple Hemiptera lineages, compensating eroded biosynthetic capabilities of primary symbionts. Bostrichidae, a family of xylophagous beetles, harbor consistently a Flavobacterial symbiont encoding exclusively the Shikimate pathway to synthesis precursors of tyrosine. However, in two families a second, closely Flavobacterial symbiont capable of recycling urea and synthesizing lysine was retained. Both symbionts exhibit high genomic syntheny and tight co-cladogenesis with the host phylogeny, indicating ancestral, ecological highly beneficial symbioses.

## Introduction

Many insects are associated with microbial partners in host-beneficial symbioses (Douglas, 2014; Feldhaar, 2011; Flórez et al., 2015; Lemoine et al., 2020). Nourishing the symbiont in specialized organs and ensuring transmission to the next generation in exchange for essential nutrients allows the host to thrive on challenging and nutritionally imbalanced diets, like plant sap, wood, or vertebrate blood (Douglas, 2009). Such stable symbiotic associations commonly experience co-evolutionary dynamics, including co-adaptation and co-speciation (Clark et al., 2000; Kikuchi et al., 2009). The continued isolation from environmental bacteria, as well as strong population bottlenecks during symbiont transmission, genetic drift, symbiont and host-level selection result in rapid genomic changes that lead to metabolic specialization of the nutritional symbiont (McCutcheon et al., 2019; Nancy A. Moran et al., 2008). The outcome is a drastically reduced symbiont genome encoding only essential pathways to sustain the symbiont’s metabolism under extensive host provisioning as well as biosynthetic pathways to supplement essential nutrients that complement the host’s metabolism (McCutcheon, 2010). Symbionts can also be lost or replaced when they are no longer needed or capable of sufficiently supporting their host’s metabolism – as genome erosion can lead to reduced efficiency in the symbionts (Bennett & Moran, 2015; Matsuura et al., 2011, 2018; Sudakaran et al., 2017).

An alternative fate of obligate symbioses is the acquisition of a second symbiont that takes over part of the original symbiont’s function or provides additional metabolic capacities to the host. This can result in multipartite symbioses with co-obligate symbionts that exhibit complementary metabolisms, e.g. dividing pathways for the essential amino acids between two symbionts (Gosalbes et al., 2008; McCutcheon & von Dohlen, 2011; Sloan & Moran, 2013) or specializing on either essential amino acids or vitamin biosynthesis, respectively (Bennett & Moran, 2013; McCutcheon & Moran, 2010; Nancy A. Moran et al., 2006). One example is the glassy-winged sharpshooter *Homalodisca vitripennis* (Hemiptera: Cicadellidae), where the β-proteobacterial symbiont *Baumannia* retains pathways for vitamins needed by the host, while the Bacteroidota symbiont *Sulcia muelleri* retains genes for the production of most essential amino acids, resulting in metabolic complementarity (Nancy A. Moran et al., 2006).

Co-obligate symbioses have so far only been functionally characterized across several lineages of Hemiptera (Koga et al., 2003; McCutcheon & Moran, 2007; McCutcheon & von Dohlen, 2011; Bublitz et al., 2019; Monnin et al., 2020). Recently, however, a dual symbiosis with two Bacteroidota was described for beetles of the family Bostrichidae (Coleoptera) (Engl et al., 2018). The family Bostrichidae (Latreille, 1802) evolved between 170 (McKenna et al., 2019) and 155 (Zhang et al., 2018) Mya and consists of phytophagous beetles. While most species are xylophagous and live and develop within twigs, branches or trunks of dead or dying trees, some species are economically important pests of wood products or stored foods, including staple roots and cereals (Jerzy Borowski & Piotr Wegrzynowicz, 2007; Niehuis, 2022). Their endosymbionts are closely related to intracellular symbionts of cockroaches and Auchenorrhyncha, i.e. *Blattabacterium* spp. and *Sulcia muelleri*, respectively, and particularly to *Candidatus* Shikimatogenerans silvanidophilus, the endosymbiont of the sawtoothed grain beetle *Oryzaephilus surinamensis* (Engl et al., 2018; Okude et al., 2017). The latter provisions tyrosine precursors to the beetle that complement the tyrosine-deficient diet of stored grain products (Kiefer et al. 2021).

Supplementation of precursors for tyrosine synthesis has been found to be important for cuticle melanisation and sclerotisation, as all of the cuticular crosslinking agents are derived from the aromatic amino acid tyrosine (Brunet, 1980; Kramer & Hopkins, 1987). Tyrosine-supplementing symbionts can inhabit the gut as in turtle ants of the genus *Cephalotes* (Duplais et al., 2021; Hu et al., 2018) or the bean weevil *Callosobruchus maculatus* (Berasategui et al., 2021), but most are located within bacteriomes, like *Candidatus* Westeberhardia cardiocondylae (Enterobacteriaceae) in the tramp ant *Cardiocondyla obscurior* (Klein et al., 2016), the γ-proteobacterial symbionts *Candidatus* Nardonella dryophthoridicola and *Candidatus* Sodalis pierantonius in weevils (Anbutsu et al., 2017; Vigneron et al., 2014), as well as the Bacteroidota endosymbiont *S. silvanidophilus* in *O. surinamensis* (Kiefer et al., 2021). The widespread occurrence of such symbioses provides evidence for this aromatic amino acid being a key nutrient for many insects to produce their strongly sclerotised and melanised exoskeleton, thereby increasing desiccation resistance and protection from predators and pathogens (Anbutsu et al., 2017; Anbutsu & Fukatsu, 2020; José de Souza et al., 2011; Kiefer et al., 2021; Vigneron et al., 2014).

Here, we set out to functionally characterize the microbial symbionts of Bostrichidae as the first dual symbiosis of insects outside of the Hemiptera. We collected 28 beetle species across five subfamilies and performed metagenome sequencing and fluorescence *in situ* hybridization. Based on the symbiont genomes and host mitochondrial genomes and nuclear markers, we reconstructed the molecular phylogenies of host and Bacteroidota symbionts. We demonstrate that (i) most or all bostrichids are associated with *Shikimatogenerans* whose genome is highly degraded, retaining only the shikimate pathway for tyrosine precursor provisioning; (ii) beetles of the genera *Lyctus* and *Dinoderus* are associated with a second Bacteroidota symbiont that encodes the capacity for lysine biosynthesis and nitrogen recycling from urea; and (iii) host and symbiont phylogenies exhibited a high degree of co-cladogenesis, indicating an ancient association that resulted in obligate mutual dependence and co-diversification. Our results shed light on the evolutionary dynamics of multipartite symbioses beyond the well-studied Hemiptera and highlight the importance of tyrosine provisioning and nitrogen recycling for the ecology of xylophagous beetles.

## Results

We collected and sequenced the metagenomes of 28 beetle species of the family Bostrichidae and supplemented our dataset with three publicly available datasets from NCBI (*Apatides fortis* mitochondrial genome [FJ613421], *Sinoxylon* sp. SIN01 mitochondrial genome [JX412742], and *Xylobiops basilaris* transcriptome [SRR2083737]) (Supplement Tables 1 and 2). The resulting 31 species covered nine tribes within five of the nine subfamilies of Bostrichidae. For thirteen of the 31 species, we were able to assemble the full and circularized genome of the Bacteroidota endosymbiont *Shikimatogenerans bostrichidophilus*, with the longest closed symbiont genome being 200,377 bp and the shortest 172,971 bp in length, and an average GC content of 15.1% (Supplement Table 2). For ten additional species, we assembled draft genomes of *Shikimatogenerans* based on multiple contigs extracted from the metagenome assemblies via taxonomic classification, GC content filtering, as well as by manually searching for tRNAs and ribosomal protein genes as well as enzymes of the shikimate pathway of Bacteroidota bacteria. For one species (*Dinoderus bifoveolatus*), we only retrieved the 16S rRNA sequence, and for another one (*Xylobiops basilaris)* only the 16S rRNA and *aroA* gene sequences of the endosymbiont. In the metagenome data of the remaining four species, we were not able to detect any sequence of Bacteroidota bacteria and the two final datasets contained only mitochondrial genomes.

As expected from a previous study (Engl et al., 2018), we found the genome of a second Bacteroidota endosymbiont, which we named *Bostrichicola ureolyticus* (see below), in some species of the subfamilies Lyctinae and Dinoderinae. In particular, we detected this co-obligate symbiont in all three *Lyctus* and two out of three *Dinoderus* species, but not in other members of the Lyctinae (*Trogoxylon impressum*) or Dinoderinae (*Rhyzopertha dominica*). The genomes of *Bostrichicola* were on average 337,500 bp in length and had an average GC content of 22.4%.

The metagenomic datasets were used to reconstruct the phylogeny of the host species (Figure 1 left and Supplement Figure 1) based on either the assembled mitochondrial genomes (Figure 1 left), or on 22 aligned and concatenated Benchmarking Universal Single-Copy Ortholog (BUSCO) genes (Waterhouse et al., 2018) found across all species (Supplement Figure 2). Both phylogenies yielded very similar results, revealing two main clades of the Bostrichidae beetles, separating the Lyctinae and Dinoderinae from the Euderinae, Apatinae and Bostrichinae. However, the two phylogenies differed in the placement of *Micrapate scabrata*, which either grouped within the Sinoxylonini as a sister clade to the Xyloperthini (BUSCO genes) or as an outgroup to the Sinoxylonini and Xyloperthini (based on the mitochondrial genomes; Supplement Figure 2).

**Figure 1:**
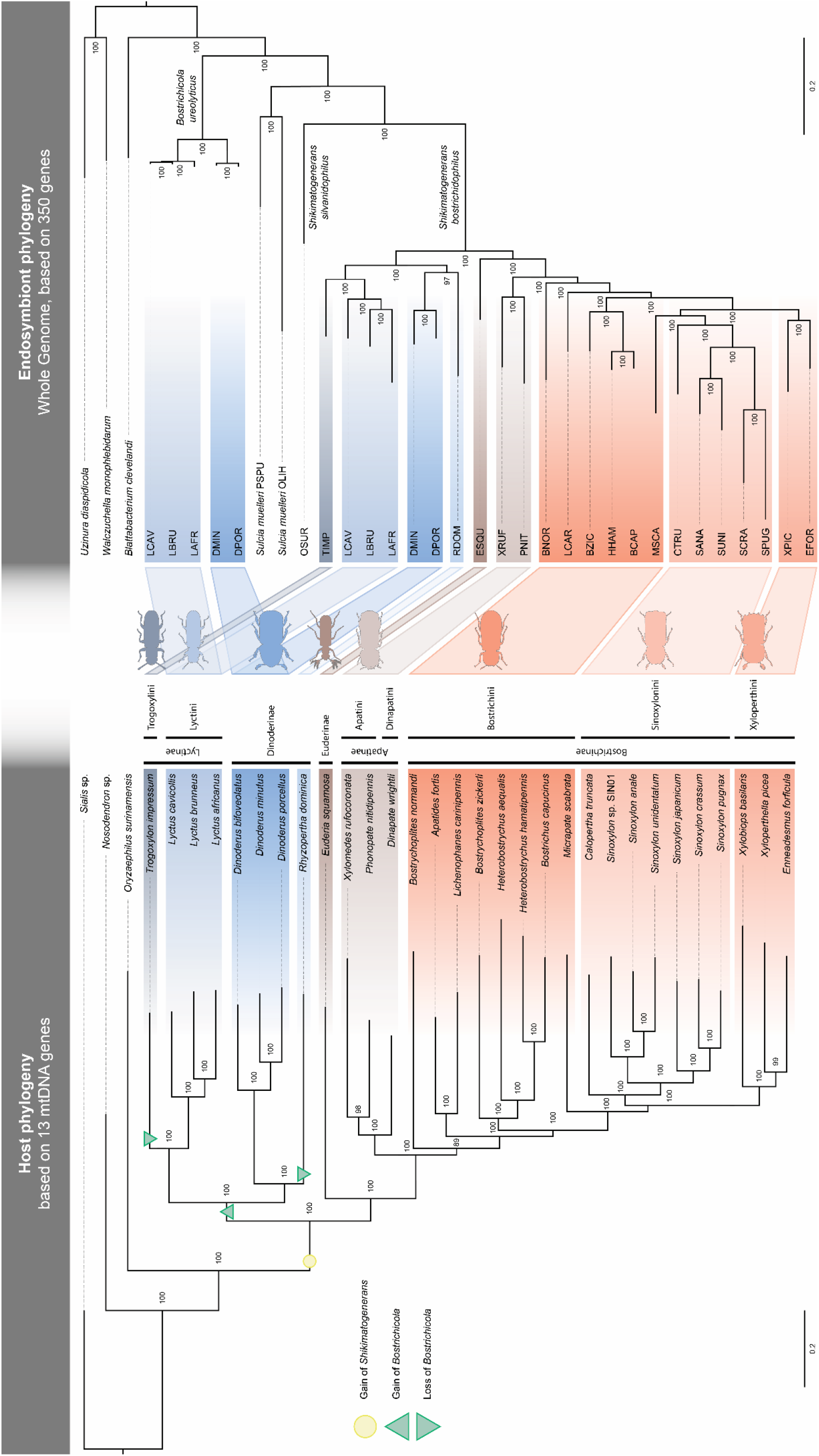
Comparison of phylogenetic relationships between Bostrichidae beetle hosts and their Bacteroidota endosymbionts. Left: Bayesian phylogeny of 30 Bostrichidae beetle species inferred from concatenated nucleotide alignment of 13 mitochondrial genes. Right: Bayesian phylogeny of Bacteroidota symbionts of Bostrichidae beetles inferred from concatenated nucleotide alignment of 350 genes. Node numbers represent posterior probabilities of Bayesian analyses. Host-symbiont associations are highlighted by connecting trapezoids between the phylogenies. Inferred, most parsimonious gain and loss events of both symbionts are indicated by circles (Shikimatogenerans) and triangles (Bostrichicola) on the host phylogeny.

Similarly, for the endosymbionts, the metagenome assemblies were used to generate a whole genome-based phylogeny (Figure 1 right). This phylogenetic reconstruction based on 350 conserved genes confirmed the monophyly of the *Shikimatogenerans* endosymbionts of Bostrichidae and Silvanidae beetles and their close relationship to other insect-associated Bacteroidota bacteria, specifically to *Blattabacterium* spp. and *Sulcia muelleri*, which had been previously reported based on 16S rRNA phylogenies (Engl et al., 2018; Hirota et al., 2017; Okude et al., 2017). The *Bostrichicola* symbionts in the *Lyctus* as well as *Dinoderus* species clustered in a distinct, more basally branching monophyletic clade within the Bacteroidota. The two clades of Bostrichidae endosymbionts were separated by the Bacteroidota symbiont *Shikimatogenerans silvanidophilus* OSUR of the sawtoothed grain beetle *Oryzaephilus surinamensis* (Silvanidae) as well as the clade of *Sulcia muelleri* endosymbionts of the Auchenorrhyncha (Hemiptera). A second phylogeny of the endosymbionts based on the 16S rRNA sequences that allowed us to include more taxa revealed a highly similar distribution within the Bacteroidota (Supplement Figure 2). The main difference between both phylogenies (Supplement Figure 2) was the placement of the endosymbiont of *Calopertha truncatula*, which formed an outgroup to endosymbionts of the Sinoxylonini and of Xyloperthini in the 16S rRNA-based phylogeny, while it was placed basally in the endosymbiont clade of Sinoxylonini within the whole genome-based phylogeny.

A comparison between the endosymbiont phylogeny based on the 350 conserved genes and the host mitochondrial phylogeny showed a high degree of co-cladogenesis (Figure 1). For *S. bostrichidophilus*, the only incongruence concerned the placement of *Micrapate scabrata* as an outgroup for the Xyloperthini and Sinoxylonini in the host phylogeny, but a grouping within the Sinoxylonini as a sister clade to the Xyloperthini in the symbiont phylogeny. However, the latter placement was also found in the BUSCO-based host phylogeny, strongly suggesting an incorrect placement in the mitochondrial phylogeny rather than a discrepancy between host and symbiont phylogenies (Supplement Figure 2). For *B. ureolyticus*, host and symbiont phylogenies were congruent on the host genus level, but the relationship of the three *Lyctus* species differed from that of their endosymbionts.

The genomes of both Bostrichidae endosymbionts were highly reduced and showed clear signs of genome erosion. Both endosymbionts retained genes involved in the cellular core processes of genetic information processing including DNA replication and repair, transcription, and translation (Supplement Figure 3). In addition, *Shikimatogenerans* encoded all of the genes of the shikimate pathway except a shikimate dehydrogenase (*aroE* [EC:1.1.1.25]) (Figure 2). Also, these genomes encoded the bifunctional *aroG*/*pheA* gene (phospho-2-dehydro-3-deoxyheptonate aldolase/chorismate mutase [EC:2.5.1.54 5.4.99.5]), capable of catalysing the Claisen rearrangement of chorismate to prephenate and the decarboxylation/dehydration of prephenate to phenylpyruvate in *Escherichia coli* (Dopheide et al., 1972).

**Figure 2:**
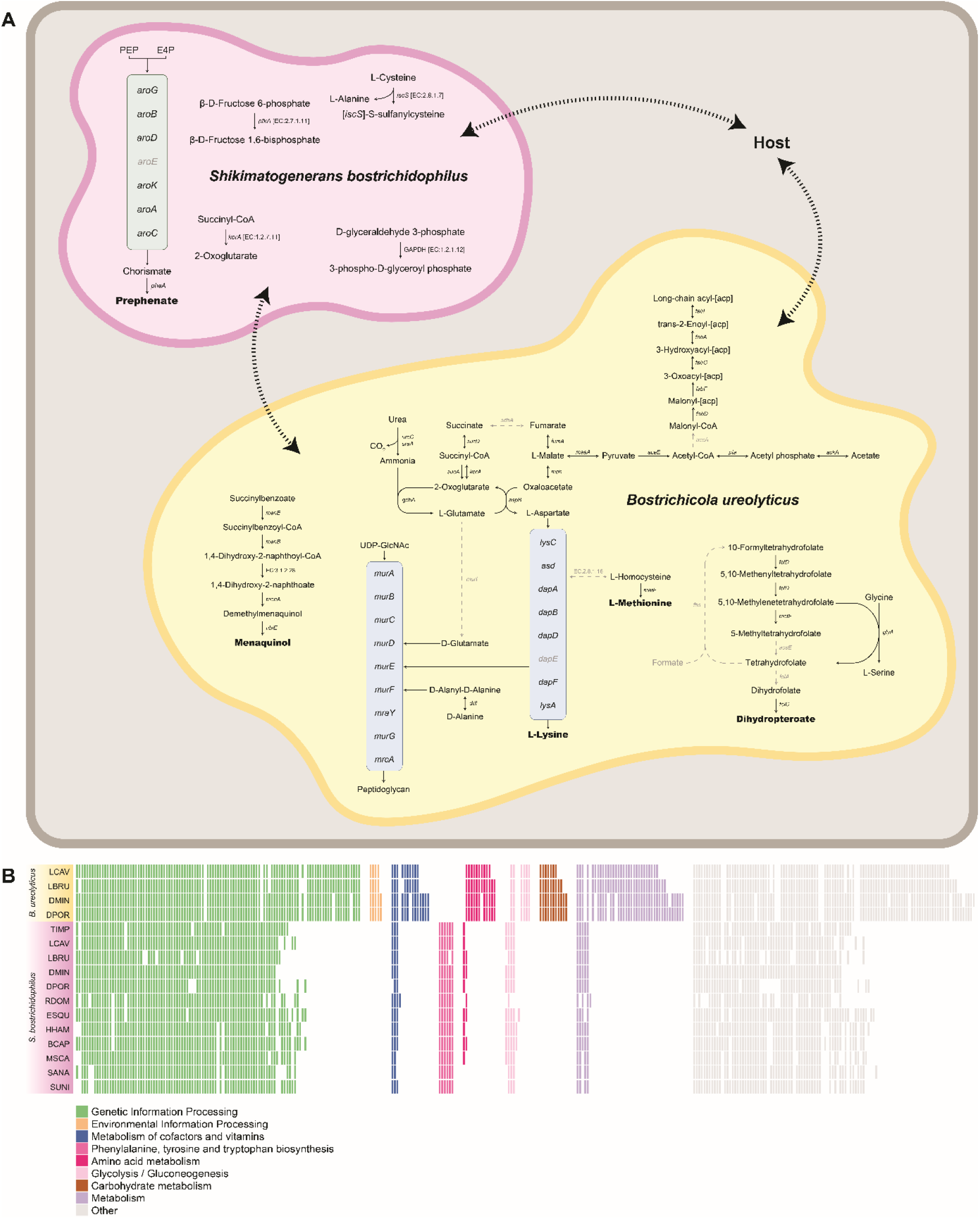
(**A**) Reconstructed metabolism of the two *Dinoderus porcellus* endosymbionts *Shikimatogenerans bostrichidophilus* DPOR *and Bostrichicola ureolyticus* DPOR, inferred from genomic data. Enzymes and arrows in grey were missing in the genome annotation. Dashed arrows indicate transport processes without annotated transporters. (**B**) Comparison of the functional gene repertoires of Bacteroidota symbionts of Bostrichid beetles that could be assembled into full continuous genomes. Coloured boxes indicate the presence, and white boxes the absence of genes in the symbiont genomes. Box colours are based on KEGG’s categories (see legend for depicted categories).

The genome of *Bostrichicola* encoded both urease α and γ subunits (*ureC* [EC:3.5.1.5]) to recycle nitrogen from urea, as well as a glutamate dehydrogenase (*gdhA* [EC:1.4.1.4]) that enables the integration of the resulting ammonium into the amino acid metabolism via glutamate (Figure 2 A). In addition, they encoded for an aspartate aminotransferase (*aspB* [EC:2.6.1.14]) to transfer the amino group from glutamate to oxaloacetate, as well as an almost complete diaminopimelate pathway to synthesize the essential amino acid lysine from aspartate. They also retained a methionine synthase to convert L-homoserine to L-methionine, a menaquinone biosynthesis pathway, and a fragmented folate biosynthesis pathway. By contrast, we did not find a single gene of the shikimate pathway to synthesize aromatic amino acids. However, the *Bostrichicola* genomes encoded for a complete fatty acid and peptidoglycan biosynthesis, albeit other cell envelope components apparently cannot be synthesized. The genomic data revealed no transporters, so it remains unknown how the symbionts exchange metabolites with the host and with each other (Figure 2 A). In addition, genes encoding signal transduction, cell surface structures, and motility were absent (Figure 2 B).

We also compared the set of genes that are not encoded in all genomes (Supplement Figure 4). For *Shikimatogenerans*, it was particularly noticeable that *mutL* (DNA mismatch repair protein) was still present in the Dinoderinae+Lyctinae symbionts but had been lost in the Euderinae+Apatinae+Bostrichinae. For *Bostrichicola*, all genes for peptidoglycan biosynthesis (*murA, murB, murC, murD, murE, murF, murG, mraY* and *mrcA*) were still encoded in the Dinoderinae, whereas the symbiont of *L. brunneus* lost *murC, murD* and *murG*, and the symbiont of *L. cavicollis* lost all of the peptidoglycan biosynthesis genes. The comparison between the full genomes of all *Shikimatogenerans* and *Bostrichicola* strains showed a high degree of synteny within, but not between, the two symbiont genera (Figure 3).

**Figure 3:**
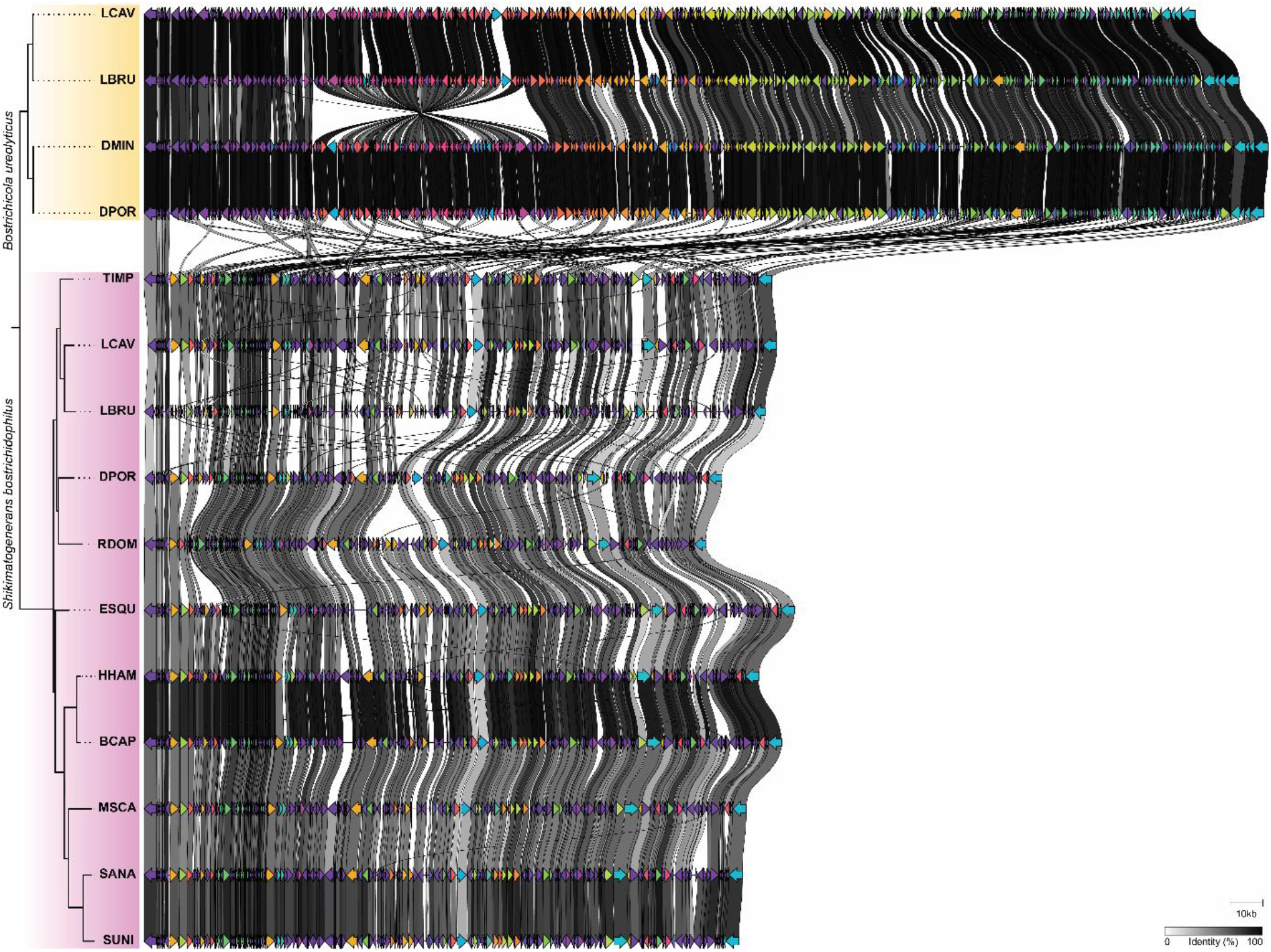
Gene order comparison between *S. bostrichidophilus* and *B. ureolyticus* genomes that could be assembled into full contiguous genomes, showing a high degree of synteny within, but not between, the two symbiont genera. Grey shades show the percentage of identity between homologous proteins from different genomes (based on amino acid sequences). The phylogenetic tree on the left is based on the symbiont phylogeny displayed in Figure 1.

Based on the close phylogenetic relationship to *Shikimatogenerans silvanidophilus* OSUR and the presence of the shikimate pathway in the highly eroded genome, we propose the name ‘*Candidatus* Shikimatogenerans bostrichidophilus*’* for the endosymbiont of Bostrichidae beetles, henceforth called *S. bostrichidophilus*. The genus name *Shikimatogenerans* refers to its ability to perform the shikimate pathway. Previous studies have shown that closely related Bacteroidota bacteria are also associated with other beetle families such as the Silvanidae and the Nosodendridae (Engl et al., 2018; Hirota et al., 2020). Thus, we propose *bostrichidophilus* as a species epithet to indicate that this symbiont clade is associated with beetles of the family Bostrichidae. As all the symbionts encode highly similar genomes, we propose to add a four-letter abbreviation of the host species to denote the strain (first letter of the host genus and first three letters of the host species epithet), e.g. *Shikimatogenerans bostrichidophilus* RDOM for the endosymbiont of *Rhyzopertha dominica*. For the second co-obligate endosymbiont found in Bostrichidae beetles of the subfamily Dinoderinae and Lyctinae, we propose the name ‘*Bostrichicola ureolyticus*’. Its genus name refers to its association with Bostrichid beetles, while *ureolyticus* refers to its metabolic potential to recycle nitrogen from urea as inferred from the genomic data. In analogy to *Shikimatogenerans*, we propose to add a four-letter abbreviation of the host species to identify the strains, e.g. *Bostrichicola ureolyticus* LBRU for the *Bostrichicola* endosymbiont of *Lyctus brunneus*.

Based on 16S rRNA fluorescence *in situ* hybridisation with eight species, the bacterial symbionts were localized intracellularly in bacteriomes in the abdomen of the host (Figure 4). The bacteriomes are located between the gut, fat body and reproductive organs, but without direct connection to any of these tissues. Bostrichid beetles of all subfamilies harbored one paired bacteriome with symbionts stained by a probe specific for members of the *Shikimatogenerans* symbiont clade (Engl et al., 2018; Kiefer et al., 2021) (Figure 4 a, e, f, g and h). In addition, species of the genera *Dinoderus* and *Lyctus* contained a second pair of bacteriomes stained by a probe specific to *B. ureolyticus* (Engl et al., 2018) (Figure 4 b, c and d). The *Bostrichicola*-harboring bacteriomes were distinct in ultrastructure, but closely adjacent to the ones containing *Shikimatogenerans*, sometimes with direct physical contact (Figure 4 b).

**Figure 4:**
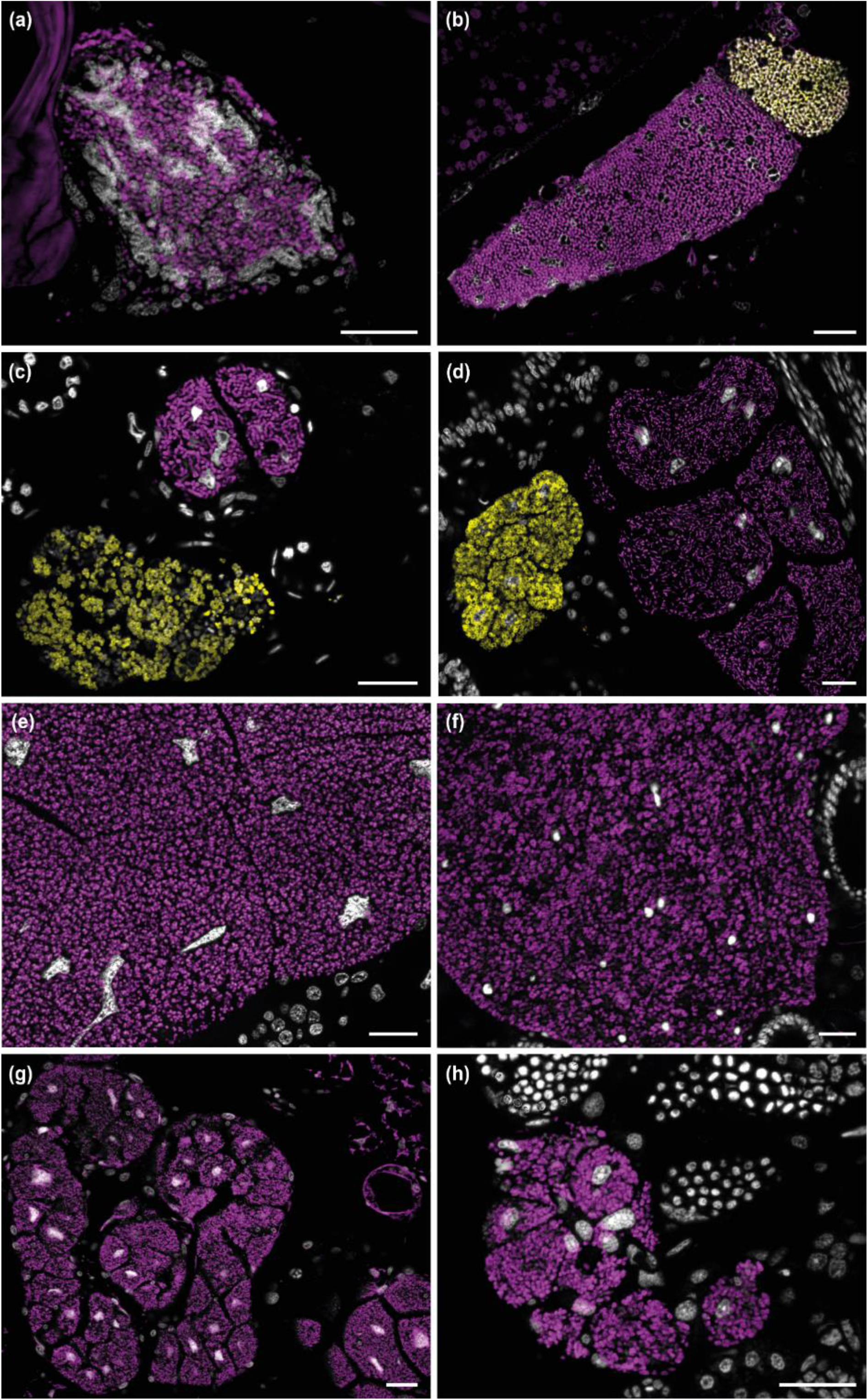
Fluorescence *in situ* hybridisation micrographs of *Shikimatogenerans* 978 *bostrichidophilus* and *Bostrichicola ureolyticus* in sections of (**a**) *Trogoxylon impressum*, (**b**) 979 *Lyctus cavicollis*, (**c**) *Dinoderus minutus*, (**d**) *Dinoderus porcellus*, (**e**) *Rhyzopertha dominica*, (**f**) 980 *Prostephanus truncatus*, (**g**) *Xyloperthella picea* and (**h**) *Sinoxylon anale*. Sections are stained 981 with a *Shikimatogenerans* specific probe (magenta), a *Bostrichicola* specific probe (yellow), 982 and DAPI targeting DNA (white). Scale bars represent 20 µm.

## Discussion

In this study, we characterized the intracellular bacterial symbionts across 29 species of auger or powderpost beetles (Coleoptera: Bostrichidae) and assessed their functional potential and co-speciation with their hosts based on comparative genomics. The functional characterization of this multipartite symbiosis enhances our understanding of the ecological relevance of microbial symbionts for a beetle family containing important wood and stored grain pest species and provides first insights into the evolutionary history and dynamics of co-obligate symbioses beyond the well-studied Hemiptera.

Based on a set of BUSCO as well as mitochondrial genomes, we reconstructed the first molecular phylogeny of Bostrichidae, after the morphological phylogeny by Liu & Schönitzer (Liu & Schönitzer, 2011). Both molecular datasets resulted in well supported phylogenies that were highly congruent and separated the Bostrichidae into two main clades: The Lyctinae and Dinoderinae grouped together, as did the Euderinae, Apatinae and Bostrichinae. The main difference was the placement of *Micrapate scabrata*, which clustered within the tribe Sinoxylonini in the BUSCO-based phylogeny, but as an outgroup to Sinoxylonini + Xyloperthini in the mitochondrial phylogeny (Supplement Figure 1). Overall, our molecular phylogenies supported the earlier morphological work, with one major exception: Liu & Schönitzer placed the Euderiinae with a single monotypic genus as a basal branch of the Bostrichidae and suggested to even place them in a separate family. In our analyses, *Euderia squamosa* robustly grouped within the Bostrichidae in a separate branch between the Lyctinae/Dinoderinae and Apatinae/Bostrichinae, confirming the affiliation of the Euderiinae to the Bostrichidae.

Given their economic importance as pests of wood or stored grains, several species of Bostrichidae beetles were intensively studied almost a century ago and found to harbor intracellular symbionts (Buchner, 1954; Gambetta, 1928; Koch, 1936; Mansour, 1934), which were recently identified as members of the insect-associated Bacteroidota (Flavobacteriaceae) clade (Engl et al., 2018; Okude et al., 2017). In our broad phylogenetic survey, we were able to detect *Shikimatogenerans bostrichidophilus* in 25 out of 29 examined Bostrichidae species. For the few species where we could not detect any symbiont (*Dinapate wrightii, Amphicerus bicaudus, Heterobostrychus aequalis*, and *Sinoxylon japanicum*), three scenarios are possible: First, it is known for several beetle taxa that the endosymbiont is degraded or lost in male beetles after metamorphosis (Fukumori et al., 2022; Janke et al., 2022; Reis et al., 2020; Vigneron et al., 2014). In cases where symbiont-provided benefits are only relevant during larval development and/or metamorphosis, males can benefit from recycling their symbionts and symbiotic organs, given that they do not transmit the symbionts to their offspring (Engl et al., 2020; Vigneron et al., 2014). Thus, as individual beetle specimens in our study may have been males, the lack of symbionts in some species may represent the absence of symbionts only in adult males rather than in all individuals. Second, in some species, aposymbiotic individuals and even populations occur in the field, due to elevated sensitivity of the symbionts to environmental stressors like heat (Dunbar et al., 2007; Koch, 1936), or possibly due to the application of certain agrochemicals like glyphosate that can eliminate symbionts encoding a sensitive *aroA* gene in the shikimate pathway (Kiefer et al. 2021). Third, *Shikimatogenerans* may have truly been lost within these species. As we had only single specimens available for the species in which we failed to detect symbionts, we cannot confidently reject any of these hypotheses. However, based on the widespread occurrence of *S. bostrichidophilus* across Bostrichidae, we can conclude that this symbiont originated from a single acquisition event at the origin of the Bostrichidae and was retained by most if not all species.

In contrast, the co-symbiont *B. ureolyticus* appears to be confined to the genera *Lyctus* and *Dinoderus*. In this case, it is unlikely that we missed the co-symbiont in other species of these two subfamilies, *T. impressum* and *R. dominica*. We had multiple individuals and life stages of *R. dominica* available as well as four specimens of *T. impressum* and never found any indication for a second bacteriome-localised symbiont, neither within our genomic datasets nor during FISH. Concordantly, previous studies (Engl et al., 2018; Okude et al., 2017) did not report on a second symbiont in *R. dominica*, while it could already be discerned based on morphologically differentiated bacteriomes in *Lyctus* and *Dinoderus* species (Engl et al., 2018; Koch, 1936).

Phylogenetic reconstructions based on either the symbiont 16S rRNA gene or 350 conservesd marker genes resulted in highly congruent phylogenies that only differed in some of the deeper splits, which were better supported in the multi-gene phylogeny than the 16S rRNA phylogeny. The endosymbiont and the host phylogenies showed a high degree of co-cladogenesis, strongly supporting single acquisition events for each symbiont and subsequent co-cladogenesis with the host, as has been found for many obligate symbionts as well as some host-specific parasites (Demastes & Hafner, 1993; N. A. Moran et al., 1993).

Within the Bostrichoidea, the Bostrichidae split from the Ptinidae between 170 (McKenna et al., 2019) and 155 Mya (Zhang et al., 2018) in the Jurassic period, which roughly coincides with the inferred age of *S. bostrichidophilus* (274-158 Mya) based on a bacterial phylogeny calibrated with estimated origins of symbiotic associations with insects (Engl et al., 2018). As closely related families within the Bostrichoidea (Dermestidae, Ptinidae) (Zhang et al., 2018) are not known to harbor bacterial endosymbionts but in some cases (Anobiinae) associate with yeast-like endosymbionts (reviewed in Martinson, 2020), *S. bostrichidophilus* was likely acquired by the ancestor of the Bostrichidae. The symbiosis with *Bostrichicola* is of somewhat more recent origin, i.e. around 100 Mya (Engl et al., 2018), and occurred in the ancestor of the Dinoderinae and Lyctinae (plus possibly Psoinae and Polycaoninae, for which we were unable to obtain specimens), and was then lost in *Rhyzopertha, Prostephanus* and *Trogoxylon*. Although the alternative scenario of two independent acquisitions in the Dinoderinae and Lyctinae seems possible, the high degree of genome synteny between the symbionts of both genera renders a single acquisition much more likely.

Both endosymbionts are characterized by extremely small, heavily eroded and A+T-biased genomes with very limited biosynthetic capabilities, akin to other strictly vertically transmitted symbionts (McCutcheon & Moran, 2012) as well as other intracellular genetic elements (Dietel et al., 2019). The genome of *S. bostrichidophilus* encodes for the shikimate pathway to synthesize precursors of aromatic amino acids. Of the seven canonical genes in the shikimate pathway (*aroG, aroB, aroD, aroE, aroK, aroA, aroC*), the *S. bostrichidophilus* genome only lacks the gene for shikimate dehydrogenase (*aroE*), which catalyses the reversible reduction of 3-dehydroshikimate to shikimate. However, the shikimate pathways of *Nardonella* EPO, the endosymbiont of the sweetpotato weevil *Euscepes postfasciatus* (Curculionidae: Cryptorhynchinae) (Anbutsu et al., 2017; Kuriwada et al., 2010) and *Carsonella ruddii*, the endosymbiont of the gall-forming psyllid *Pachypsylla venusta* (Aphalaridae: Pachypsyllinae) (Sloan et al., 2014), as well as the closely related *S. silvanidophilus* OSUR in the sawtoothed grain beetle *O. surinamensis* also lack *aroE*, but remain functional (Anbutsu et al., 2017; Kiefer et al., 2021), indicating that the function of *aroE* is taken over by other enzymes of either host or endosymbiont origin. Hence, *S. bostrichidophilus* is inferred to transform phosphoenolpyruvate (PEP) and erythrose-4-phosphate (E4P) to prephenate/chorismate via the shikimate pathway (Herrmann & Weaver, 1999; Mir et al., 2015), which can then be converted by the host to the aromatic amino acid tyrosine (Arakane et al., 2009). As all of the cuticular crosslinking agents as well as precursors for black and brown pigments (melanins) are derived from tyrosine (Brunet, 1980; Kramer & Hopkins, 1987), it constitutes the key metabolite in cuticle synthesis, melanisation and sclerotisation, thereby strongly affecting the physicochemical properties of the cuticle (Hackman, 1974). Concordantly, symbiont-mediated supplementation of tyrosine precursors enables the production of a thicker, stronger, and darker cuticle and thereby enhances protection against desiccation, predation, and infection by entomopathogens (Anbutsu et al., 2017; Anbutsu & Fukatsu, 2020; Engl et al., 2018; Kanyile et al., 2022).

The reduced genome of the co-obligate symbiont *B. ureolyticus* encodes the genes for urea recycling and the diaminopimelate pathway to synthesize lysine. In addition, *B. ureolyticus* retained partial pathways to convert intermediates of the lysine biosynthesis into methionine, folate and menaquinone (Douglas, 2009; Wu et al., 2006), and it can synthesize some components of the cell envelope (fatty acids and peptidoglycan). However, several of these pathways exhibit differential erosion between the *Lyctus* and *Dinoderus* symbionts, but even between the symbionts of one of the genera (Supplemental Fig. S4). Especially the pathways for synthesis of the cell envelopes are intriguing, as cell wall synthesis has been demonstrated to be complemented and controlled by host encoded genes and inter-symbiont exchange of metabolites in hemipteran symbionts (Bublitz et al., 2019; Smith et al., 2021, 2022).

Nitrogen recycling is well documented within some Bacteroidota endosymbionts (Hansen et al., 2020; Rosas-Pérez et al., 2014; Sabree et al., 2009) and can be an important benefit for insects developing in nitrogen-limited diets (Hoadley et al., 1998; Jerzy Borowski & Piotr Wegrzynowicz, 2007; Niehuis, 2022; Oke et al., 1990; Souci et al., 2009). In Bostrichid beetles, the recycling of urea as a source of amino groups is likely important for the formation of tyrosine, but also other amino acids and amino acid-derived components of the cuticle like N-acetyl-glucosamine, the monomer of chitin (Duplais et al., 2021). However, why *B. ureolyticus* retained a lysine biosynthesis pathway is less clear. Lysine constitutes an important amino acid of cuticular proteins, as its ε-amino group represents an anchor point for cross linking (Suderman et al., 2010). In addition, grain diets, but also staple roots are specifically limited in lysine (Juliano, 1999; Torbatinejad et al., 2005), so symbiont-mediated lysine supplementation could be an important benefit for the stored product pest beetles of the genus *Dinoderus*, but also other species of the Dinoderinae subfamily. However, a deeper understanding of why certain genera of Bostrichidae benefit from such provisioning and thus retain *Bostrichicola* while others do not, is currently hampered by the scarcity of information on the ecology of most Bostrichidae (Liu & Schönitzer, 2011).

Based on our phylogenetic analyses, *Shikimatogenerans* and *Bostrichicola* are derived from the same ancestor as *Blattabacterium* spp., *Walczuchella monophlebidarum* and *Uzinura diaspidicola*, but then diverged and evolved different functional specializations (Engl et al., 2018; Hirota et al., 2020; Kiefer et al., 2021; Sabree et al., 2013). Interestingly, *S. silvanidophilus* OSUR (Kiefer et al., 2021) *–* the sister taxon of *S. bostrichidophilus* – retained the shikimate pathway as well as one urease subunit putatively involved in nitrogen recycling (Kiefer et al., 2021) - and there is evidence for nitrogen recycling in *Blattabacterium* (Hansen et al., 2020; Ló Pez-Sánchez et al., 2009; Sabree et al., 2009) and *Walczuchella* (Rosas-Pérez et al., 2014), so urea catabolism seems to be a widespread and possibly ancestral benefit provided by Bacteroidota symbionts of insects.

Beneficial associations with two metabolically complementary symbionts synthesizing essential amino acids and vitamins have thus far only been functionally characterized for multiple different lineages of plant sap-feeding Hemiptera (Manzano-Marín et al., 2018; McCutcheon et al., 2009; McCutcheon & Moran, 2007, 2010; McCutcheon & von Dohlen, 2011; Nancy A. Moran, 2006; Snyder & Rio, 2015; Wu et al., 2006). In these cases, however, the co-obligate symbionts usually originated from different classes or phyla, with the exception of some associations between Sternorrhyncha and two co-obligate γ-proteobacterial symbionts (von Dohlen et al., 2017). To our knowledge, the Bostrichidae are thus far unique in containing species that harbor two Bacteroidota endosymbionts, which are closely related but diverged to metabolically complementary symbionts. This finding highlights the versatile nature of symbioses with Bacteroidota bacteria that are emerging as widespread beneficial symbionts across at least four beetle families (Silvanidae: (Engl et al., 2018; Hirota et al., 2017), Coccinellidae: (Hurst et al., 1996; Hurst, Bandi, et al., 1999), Nosodendridae (Hirota et al., 2020), and Bostrichidae: this study and Engl et al. 2018) as well as at least two other insect orders (Bandi et al., 1995; Engl et al., 2018; Nancy A. Moran et al., 2006). The repeated independent acquisitions of these specific clades of Bacteroidota symbionts suggest that these bacteria were once specialized in establishing lasting infections in insects, akin to *Wolbachia* (Kiefer et al., 2022) or *Sodalis* (McCutcheon et al., 2019). Considering that a basal clade of Bacteroidota endosymbionts are male-killing endosymbionts in different ladybird beetles (Hurst et al., 1996; Hurst, Bandi, et al., 1999; Hurst, Jiggins, et al., 1999), it is conceivable that the ancestral success of Bacteroidota symbionts relied on reproductive manipulation to spread in insect populations. Later, some of these originally parasitic associations may have evolved into beneficial symbioses due to the symbionts’ capacity to provision limiting nutrients, in analogy to the vitamin-provisioning *Wolbachia* symbionts in bedbugs (Hosokawa et al. 2010). Given the sometimes severe fitness consequences of reproductive manipulation on the host and the ensuing evolutionary arms race between host and parasite (Charlat, Hornett, et al., 2007; Charlat, Reuter, et al., 2007), the parasitic interactions may have been evolutionarily labile. By contrast, interactions that evolved towards mutualism likely remained long-term stable, which may explain the bias towards beneficial interactions observed in extant insect-associated Bacteroidota.

## Material and Methods

### Insect collection

Specimens of 28 species were collected or provided by experts in the field from Germany, the Czech Republic, Yemen, the United Arabic Emirates, the United States of America, Japan, and New Zealand (Supplement Table 1), in compliance with the Nagoya protocol. In addition, three publicly available data sets of Bostrichid beetles were retrieved from NCBI (SRR2083737, FJ613421 and JX412742).

### Symbiont genome sequencing, assembly, and annotation

Total DNA was isolated using the Epicentre MasterPure™ Complete DNA and RNA Purification Kit (Illumina Inc., Madison, WI, USA) including RNase digestion, or the QIAGEN Genomic-tip kit using 20/G columns (Qiagen, Hilden, Germany). Short-read library preparation and sequencing were performed at the Max-Planck-Genome-Centre Cologne, Germany (SRR19201352 - SRR19201388) on a HiSeq3000 Sequencing System (Illumina Inc., Madison, WI, USA), or at CeGaT on a HiSeq2500 Sequencing System (Tübingen, Germany) or a MiSeq (Illumina Inc., Madison, WI, USA) of AIST Japan (DRR414867). Adaptor and quality trimming was performed with Trimmomatic (Bolger et al., 2014).

Long-read sequencing for *D. porcellus* (SRR19201386 and SRR19201352) and *L. brunneus* (SRR19201357) was performed on a MinION Mk1B Sequencing System (Oxford Nanopore Technologies (ONT), Oxford, UK). Upon receipt of flowcells, and again immediately before sequencing, the number of active pores on flowcells was measured using the MinKNOW software (v18.12.9 and 19.05.0, ONT, Oxford, UK). Flowcells were replaced into their packaging, sealed with parafilm and tape, and stored at 4°C until use. Library preparation was performed with the Ligation Sequencing Kit (SQK-LSK109, ONT, Oxford, UK) and completed libraries were loaded on a flowcell (FLO-MIN106D, ONT, Oxford, UK) following the manufacturer’s instructions. PacBio long-read sequencing of *D. porcellus* (SRR19201385) was performed at the Max-Planck-Genome-Centre Cologne, Germany on a Sequel II system (PacBio, Menlo Park, CA, USA).

Quality-controlled long reads were taxonomy-filtered using a custom-made kraken2 database (Wood et al., 2019; Wood & Salzberg, 2014) containing the publicly available genomes of Bacteroidota bacteria to extract beetle-associated Bacteroidota sequences using the supercomputer Mogon of the Johannes Gutenberg-University (Mainz, Germany). Assembly of Illumina reads and additional hybrid assemblies with long-read libraries were performed using SPAdes (v3.15.0) with default settings (Bankevich et al., 2015). The resulting contigs were binned using BusyBee Web (Laczny et al., 2017), and screened for GC content and taxonomic identity to Bacteroidota bacteria. The extracted contigs were de novo assembled in Geneious Prime 2019 (v2019.1.3, https://www.geneious.com). The resulting contigs were then automatically annotated with PROKKA (Seemann, 2014) using the app Annotate Assembly and Re-annotate Genomes (v1.14.5) in KBase (Arkin et al., 2018). Synteny analysis of complete endosymbiont genomes was performed using Clinker with default settings (Gilchrist & Chooi, 2021).

### Fluorescence *in situ* hybridisation

Endosymbionts of *D. minutus, D. porcellus, L. cavicollis, P. truncates, R. dominica, S. anale, T. impressum* and *X. picea* were localised by fluorescence *in situ* hybridisation (FISH) on semi-thin sections of adult beetles, targeting the 16S rRNA sequence. Adult beetles were fixated in 80% tertiary butanol (Roth, Karlsruhe, Germany), 3.7% paraformaldehyde (Roth, Karlsruhe, Germany) and 3.7% glacial acetic acid (Sigma-Aldrich, Taufkirchen, Germany) for 2 hours, followed by post-fixation in alcoholic formaldehyde (3.7% paraformaldehyde and 80% tertiary butanol). After dehydration, the specimens were embedded in Technovit 8100 (Kulzer, Germany)^100^ and cut into 8 µm sagittal sections using a Leica HistoCore AUTOCUT R microtome (Leica, Wetzlar, Germany) equipped with glass knives. The obtained sections were mounted on silanised glass slides. For FSIH, each slide was covered with 100 µL of hybridization mix, consisting of hybridization buffer (0.9 M NaCl, 0.02 M Tris/HCl pH 8.0, 0.01% SDS; Roth, Germany) and 0.5 µM of the *Shikimatogenerans bostrichidophilus*-specific probe (5’-CTTCCTACACGCGAAATAG-3’; Engl et al. 2018) labelled with Cy5, as well as the *Bostrichicola ureolyticus*-specific probe (5’-TACTCGATGGCAATTAACAAC-3’; Engl et al. 2018) labelled with Cy3. DAPI (0.5 µg/mL) was included as a general counterstain for DNA. Slides were covered with glass cover slips and incubated in a humid chamber at 50°C overnight. After washing and incubating them for 20 minutes at 50°C in wash buffer (0.1 M NaCl, 0.02 M Tris/HCl, 5 mM EDTA, 0.01% SDS), they were washed in deionized water for 20 minutes, dried and mounted with Vectashield (Vector Laboratories, Burlingame, CA, USA). The sections were observed under a Zeiss AxioImager.Z2 equipped with an Apotome.2 (Zeiss, Jena, Germany) and illuminated by a SOLA Light Engine (Lumencor, Beaverton, OR, USA).

### Phylogenetic analyses

We generated phylogenetic trees based on the metagenome data generated from our Bostrichid taxa (SRR19201352 - SRR19201388) as well as three Bostrichidae shotgun sequencing datasets available on NCBI (SRR2083737, FJ613421 and JX412742).

A phylogenetic tree of the mitochondrial genes of the hosts was reconstructed by assembling the mitochondrial genome using NOVOPlasty (Dierckxsens et al., 2017) and MitoZ (Meng et al., 2019) and afterwards annotating them with Mitos (Bernt et al., 2013) (http://mitos.bioinf.uni-leipzig.de/index.py). Subsequently, 13 mitochondrial genes were translated and aligned using MUSCLE (Edgar, 2004) (v3.8.425) as implemented in Geneious Prime 2019 (v2019.1.3, https://www.geneious.com). Additionally, we generated a second (codon-based) nucleotide alignment based on Benchmarking Universal Single-Copy Orthologs (BUSCO) using a custom pipeline (Waterhouse et al., 2018) to extract the genes from the metagenome datasets. BUSCO analysis was performed for each dataset using the insecta_odb10 database (1,658 genes) to extract BUSCO genes that were found across all species (Shin et al., 2021). The corresponding nucleotide sequences were then extracted and aligned with MAFFT (Katoh & Standley, 2013) with --auto and default options. Gaps in the resulting alignment were then trimmed from the alignment using trimAl (v1.2), accepting 5% gaps for each position (Capella-Gutierrez et al., 2009). Afterwards, the aligned nucleotide sequences for each taxon were concatenated.

For the phylogenetic analyses of the intracellular symbionts of Bostrichid beetles, coding sequences were extracted from the genomes, aligned based on the nucleotide sequence with MAFFT (Katoh & Standley, 2013), and concatenated in Geneious Prime 2019 (v2019.1.3, https://www.geneious.com). Additionally, beetle symbiont 16S rRNA sequences were aligned to representative Bacteroidota 16S rRNA sequences obtained from the NCBI database, using the SILVA algorithm (Engl et al., 2018; Quast et al., 2013; Yilmaz et al., 2014). Since complete genomes were not available for some of the species, the 16S rRNA alignment allowed us to incorporate a larger number of species in the phylogenetic analysis, albeit at a lower resolution due to the limited amount of information contained in this single gene.

Phylogenetic reconstructions for all alignments were done by Bayesian inference applying a GTR+G+I model using MrBayes (v3.2.7) (Abadi et al., 2019; J. P. Huelsenbeck et al., 2001; John P Huelsenbeck & Ronquist, 2001; Ronquist & Huelsenbeck, 2003). The analysis ran for 10,000,000 generations with a “Burnin” of 25% and tree sampling every 1,000 generations. We confirmed that the standard deviation of split frequencies converged to < 0.01. The obtained trees were visualized using FigTree (v1.4.4, http://tree.bio.ed.ac.uk/software/figtree/).

## Data Accessibility Statement

Sequencing libraries and the assembled genome of the Bostrichid symbionts (*Shimatogenerans bostrichidophilus* and *Bostrichicola ureolyticus*) were uploaded to the NCBI and DDBJ Sequence Read Archives (see Supplement Table 1 for accession numbers) and GenBank (see Supplement Table 2 for accession numbers). Alignments used for all phylogenetic analyses and tree and vector graphic files of all phylogenies as well as the annotated X. basilaris mitochondrial genome are available on the data repository of the Max-Planck-society Edmond (Engl et al., 2022).

## Acknowledgments

We thank Philipp-Martin Bauer, Hans-Georg Folz, Cornel Adler, Rudy Plarre, Rich Leschen, Michael Eifler, Miguel Diaz, Ryutaro Iwata, Hiroki Watanabe, Tsuyoshi Yoshimura, Akihiro Miyanoshita and Shigeru Kuratani for providing Bostrichid specimens. We thank Benjamin Weiss for technical assistance in histology, Bruno Hüttel and the Max Planck-Genome-Centre Cologne (http://mpgc.mpipz.mpg.de/home/) for performing library preparation and sequencing of most samples in this study, Minoru Moriyama for support on bioinformatic analyses, Yu Okamura for help with the BUSCO pipeline as well as the Johannes Gutenberg-University Mainz for computation time granted on the supercomputer ‘MOGON’, and Christian Meesters for administrative assistance on ‘MOGON’. M.K. and T.E. acknowledge funding from the Max Planck Society, and further financial support of the Johannes Gutenberg-University Mainz (intramural funding to T.E.), as well as a Consolidator Grant of the European Research Council (ERC CoG 819585 *“*SYMBeetle*”* to M.K.). T.F. was supported by the Japan Science and Technology Agency (JST) ERATO Grant Number JPMJER1902.

## Contributions

J.S.T.K., T.E. and M.K. designed the project, and J.S.T.K., E.B., G.O., T.F. and T.E. sequenced and assembled the symbiont genomes. J.S.T.K. and E.B. annotated the genomes and performed symbiont genomic analysis and J.S.T.K. and T.E. performed phylogenetic analyses. J.S.T.K. and T.E. wrote the initial manuscript, with input from M.K. All authors read and commented on the manuscript.

