## Supplementary material for "Cuticle supplementation and nitrogen recycling by a dual bacterial symbiosis in a family of xylophagous beetles (Coleoptera: Bostrichidae)"

**Table S1:** List of collected Bostrichidae specimens. JKI = Julius Kühn-Institute / Federal Research Centre for Cultivated Plants; BAM = Federal Institute for Materials Research and Testing.

| Family | Subfamily | Tribe | Genus | Species | Collected | Location | Source | Library | BUSCO | mtDNA | Coverage |
|---|---|---|---|---|---|---|---|---|---|---|---|
| Bostrichidae | Lyctinae | Trogoxylini | *Trogoxylon* | *impressum* | 21.06.2020 | Engelstadt, Germany | Hans-Georg Folz | SRR19201388 | 82% | ON707259 | 7,034 |
| | | Lyctini | *Lyctus* | *africanus* | 18.03.2016 | Tokyo, Japan | Ryutaro Iwata | SRR19201387 | 66% | ON707246 | 2,914 |
| | | | | *brunneus* | 2015 | lab-culture; BAM, Berlin, Germany | Rudy Plarre | SRR19201365, SRR19201357 | 72% | ON707247 | 7,068 |
| | | | | *cavicollis* | June 2020 | Engelstadt, Germany | Hans-Georg Folz | SRR19201376 | 89% | ON707249 | 8,533 |
| | Dinoderinae | | *Dinoderus* | *bifoveolatus* | 2014 | lab-culture; JKI, Berlin, Germany | Cornel Adler | SRR19201356 | 54% | ON707238 | 752 |
| | | | | *minutus* | 12.04.2016 | Oita, Japan | Hiroki Watanabe | SRR19201354, SRR19201355 | 30% | ON707239 | 5,902 |
| | | | | *porcellus* | 2014 | lab-culture; JKI, Berlin, Germany | Cornel Adler | SRR19201352, SRR19201353, SRR19201382- | 63% | ON707240 | 9,216 |
| | | | *Rhyzopertha* | *dominica* | 2014 | lab-culture; JKI, Berlin, Germany | Cornel Adler | SRR19201381, SRR19201380 | 77% | ON707253 | 3,455 |
| | | | | | 1990 | lab-culture; Japan | Akihiro Miyanoshita | DRR414867 | | | |
| | Euderiinae | | *Euderia* | *squamosa* | 2020 | Auckland, New Zealand | Rich Leschen | SRR19201379 | 55% | ON707243 | 1,421 |
| | Apatinae | Apatini | *Phonapate* | *nitidipennis* | 29.06.1992 | Mabar, Yemen | Michael Eifler | SRR19201378 | 1% | ON707252 | 2,806 |
| | | | *Xylomedes* | *rufocoronata* | 22.03.2006 | Emirate of Sharjah, UAE | Michael Eifler | SRR19201377 | 97% | ON707261 | 27,36 |
| | | Dinapatini | *Dinapate* | *wrightii* | July 1994 | Riverside county, CA, USA | BioQuip | SRR19201375 | 3% | ON707241 | 893 |
| | Bostrichinae | Bostrichini | *Amphicerus* | *bicaudus* | 15.06.1988 | Lubbock county, TX, USA | BioQuip | SRR19201374 | 4% | - | - |
| | | | *Apatides* | *fortis* | - | - | NCBI | FJ613421 | - | - | - |
| | | | *Bostrichus* | *capucinus* | May 2020 | Ostrovačice, Czech Republic | Miguel Diaz | SRR19201373 | 98% | ON707234 | 81,589 |
| | | | *Bostrychoplites* | *normandi* | 01.08.1992 | ʿAmrān, Yemen | Michael Eifler | SRR19201371 | 1% | ON707235 | 399 |
| | | | | *zickeli* | 15.09.2001 | Al Kawd, Yemen | Michael Eifler | SRR19201372 | 72% | ON707236 | 4,253 |
| | | | *Heterobostrychus* | *aequalis* | 12.09.2016 | Kyoto, Japan | Tsuyoshi Yoshimura | SRR19201370 | 97% | ON707244 | 3,984 |
| | | | | *hamatipennis* | 01.05.2016 | Mannheim, Germany | Philipp-Martin Bauer | SRR19201369 | 70% | ON707245 | 3,47 |
| | | | *Lichenophanes* | *carinipennis* | 08.08.2020 | Kobe, Japan | Shigeru Kuratani | SRR19201368 | 64% | ON707248 | 5,899 |
| | | | *Micrapate* | *scabrata* | 01.07.2018 | University of Hohenheim, Germany | Philipp-Martin Bauer | SRR19201367 | 50% | ON707250 | 6,625 |
| | | Xyloperthini | *Enneadesmus* | *forficula* | 31.01.2006 | Emirate of Sharjah, UAE | Michael Eifler | SRR19201366 | 86% | ON707242 | 1,01 |
| | | | *Xylobiops* | *basilaris* | - | - | NCBI | SRR2083737 | 72% | - | 103,675 |
| | | | *Xyloperthella* | *picea* | 15.06.2002 | Taiz, Yemen | Michael Eifler | SRR19201364 | 55% | ON707260 | 6,599 |
| | | Sinoxylonini | *Calopertha* | *truncatula* | 12.07.2001 | Kawd al-ʿAbadil, Yemen | Michael Eifler | SRR19201363 | 97% | ON707237 | 2,116 |
| | | | *Sinoxylon* | *anale* | 01.04.2016 | Mannheim, Germany | Philipp-Martin Bauer | SRR19201362 | 69% | ON707254 | 4,303 |
| | | | | *crassum* | 01.07.2018 | University of Hohenheim, Germany | Philipp-Martin Bauer | SRR19201361 | 71% | ON707255 | 8,405 |
| | | | | *japanicum* | 26.04.2020 | Yokohama, Japan | Ryutaro Iwata | SRR19201360 | 83% | ON707256 | 1,681 |
| | | | | *pugnax* | 05.06.2005 | Emirate of Fujairah, UAE | Michael Eifler | SRR19201359 | 21% | ON707257 | 384 |
| | | | | *unidentatum* | 01.07.2018 | University of Hohenheim, Germany | Philipp-Martin Bauer | SRR19201358 | 75% | ON707258 | 6,251 |
| | | | | *sp.* | - | - | NCBI | JX412742 | - | - | - |

**Table S2:** General features of the symbiont genomes based on annotations with Prokka. LSU: Large subunit ribosomal protein, SSU: Small subunit ribosomal proteins.

| Bacterium | Strain | Accession | Contigs | Genome Size | GC content | Coverage | Predicted proteins | Transfer RNAs | Ribosomal RNAs | SSU | LSU |
|---|---|---|---|---|---|---|---|---|---|---|---|
| | LAFR | JAMYEW000000000 | 171 | 263,976 bp | 22,6% | 338 | 277 | 28 | 29 | 14 | 15 |
| | LBRU | CP100318 | 1 | 337,336 bp | 22,1% | 55 | 324 | 20 | 44 | 17 | 27 |
| *Bostrichicola ureolyticus* | LCAV | CP100320 | 1 | 323,829 bp | 22,1% | 693 | 317 | 20 | 44 | 17 | 27 |
| | DMIN | CP100321 | 1 | 346,375 bp | 22,6% | 997 | 336 | 21 | 44 | 17 | 27 |
| | DPOR | CP100319 | 1 | 344,793 bp | 22,8% | 334 | 334 | 20 | 42 | 16 | 26 |
| | TIMP | CP099825 | 1 | 193,392 bp | 13,2% | 85 | 193 | 21 | 36 | 17 | 19 |
| | LAFR | JAMYEV000000000 | 18 | 187,750 bp | 16,7% | 115 | 162 | 24 | 33 | 15 | 18 |
| | LBRU | CP099821 | 1 | 191,425 bp | 13,5% | 17 | 222 | 22 | 35 | 16 | 19 |
| | LCAV | CP099823 | 1 | 194,907 bp | 13,2% | 450 | 191 | 21 | 34 | 16 | 18 |
| | DMIN | CP099824 | 1 | 194.383 bp | 13,9% | 128 | 197 | 21 | 35 | 15 | 20 |
| | DPOR | CP099822 | 1 | 177,818 bp | 13,7% | 242 | 178 | 18 | 35 | 17 | 18 |
| | RDOM | CP099826 | 1 | 172,971 bp | 13,3% | 1,751 | 168 | 22 | 37 | 17 | 20 |
| | ESQU | CP099827 | 1 | 200,377 bp | 15,1% | 88 | 200 | 20 | 38 | 18 | 20 |
| | PNIT | CP099828 | 1 | 163,888 bp | 15,4% | 1,159 | 175 | 24 | 30 | 13 | 17 |
| | XRUF | JAMYEX000000000 | 2 | 131,783 bp | 14,7% | 27 | 129 | 17 | 33 | 15 | 18 |
| | BCAP | CP099829 | 1 | 196,478 bp | 17,1% | 1,542 | 191 | 21 | 41 | 18 | 23 |
| *Shikimatogenerans bostrichidophilus* | BNOR | JAMYEY000000000 | 49 | 117,215 bp | 15,3% | 396 | 111 | 13 | 28 | 13 | 15 |
| | BZIC | JAMYEZ000000000 | 54 | 154,309 bp | 14,5% | 47 | 154 | 15 | 25 | 11 | 14 |
| | HHAM | CP099830 | 1 | 189,381 bp | 17,3% | 184 | 187 | 19 | 40 | 18 | 22 |
| | LCAR | ON922557-ON922562 | 6 | 50,604 bp | 17,6% | 282 | 64 | 4 | 25 | 12 | 13 |
| | MSCA | CP099831 | 1 | 185,513 bp | 16,6% | 151 | 182 | 21 | 37 | 16 | 21 |
| | EFOR | JAMYFA000000000 | 13 | 164,672 bp | 15,6% | 281 | 161 | 21 | 37 | 17 | 20 |
| | XPIC | JAMYFC000000000 | 12 | 170,362 bp | 13,2% | 281 | 163 | 27 | 37 | 17 | 20 |
| | CTRU | JAMYFB000000000 | 38 | 128,631 bp | 16,5% | 55 | 128 | 20 | 36 | 15 | 21 |
| | SANA | CP099832 | 1 | 184,356 bp | 15,5% | 199 | 181 | 20 | 38 | 17 | 21 |
| | SCRA | ON922555-ON922556 | 2 | 28,977 bp | 19,8% | 463 | 26 | 7 | 7 | 3 | 4 |
| | SPUG | ON922537-ON922554 | 18 | 45,556 bp | 19,8% | 27 | 50 | 3 | 6 | 4 | 2 |
| | SUNI | CP099833 | 1 | 183,323 bp | 17,0% | 587 | 184 | 20 | 38 | 17 | 21 |

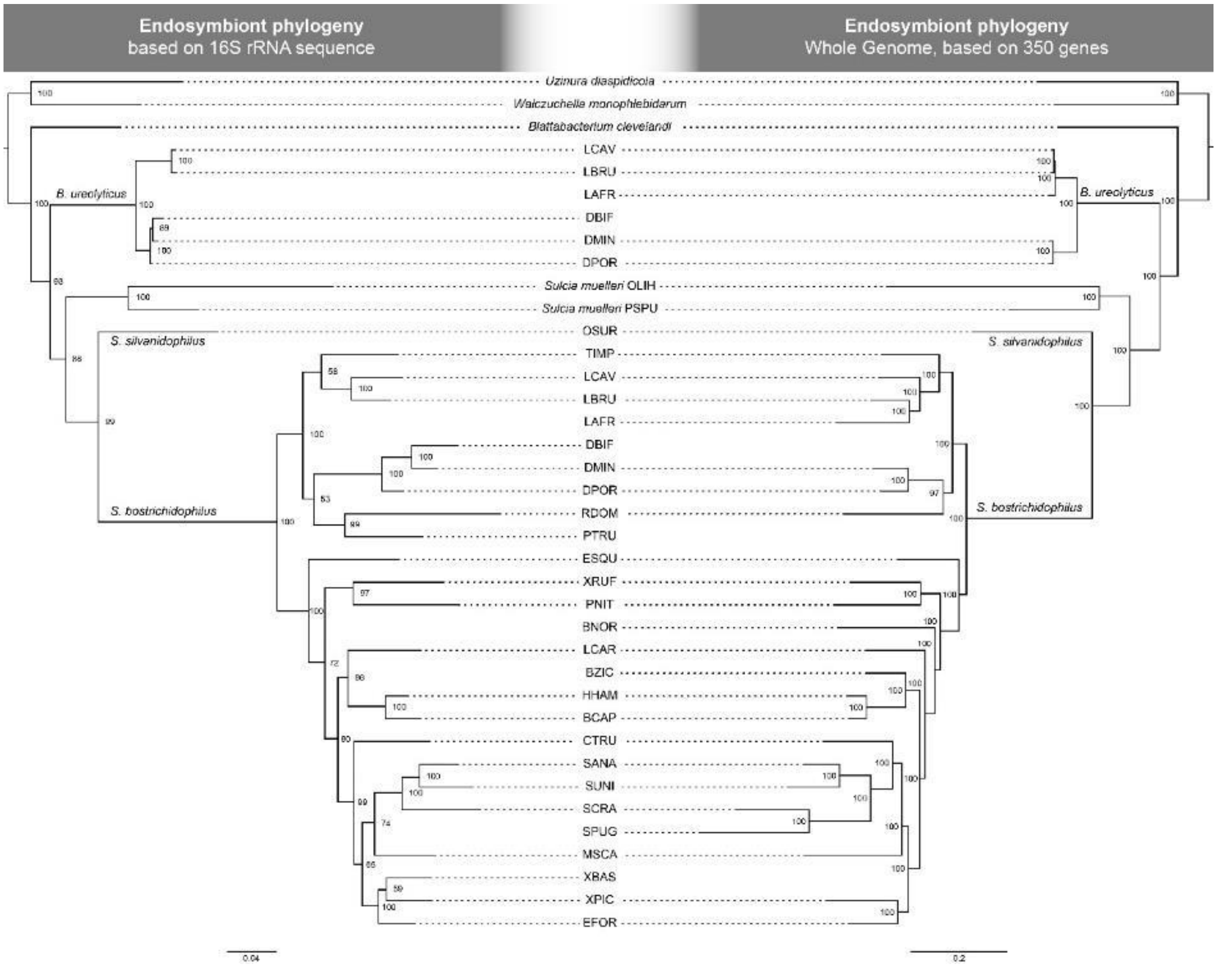


**Figure S1:** Comparison of symbiont phylogenies based on the 16S rRNA gene alone (left), and 350 genes conserved across at least two genomes (right). Node numbers represent posterior probabilities of Bayesian analyses.

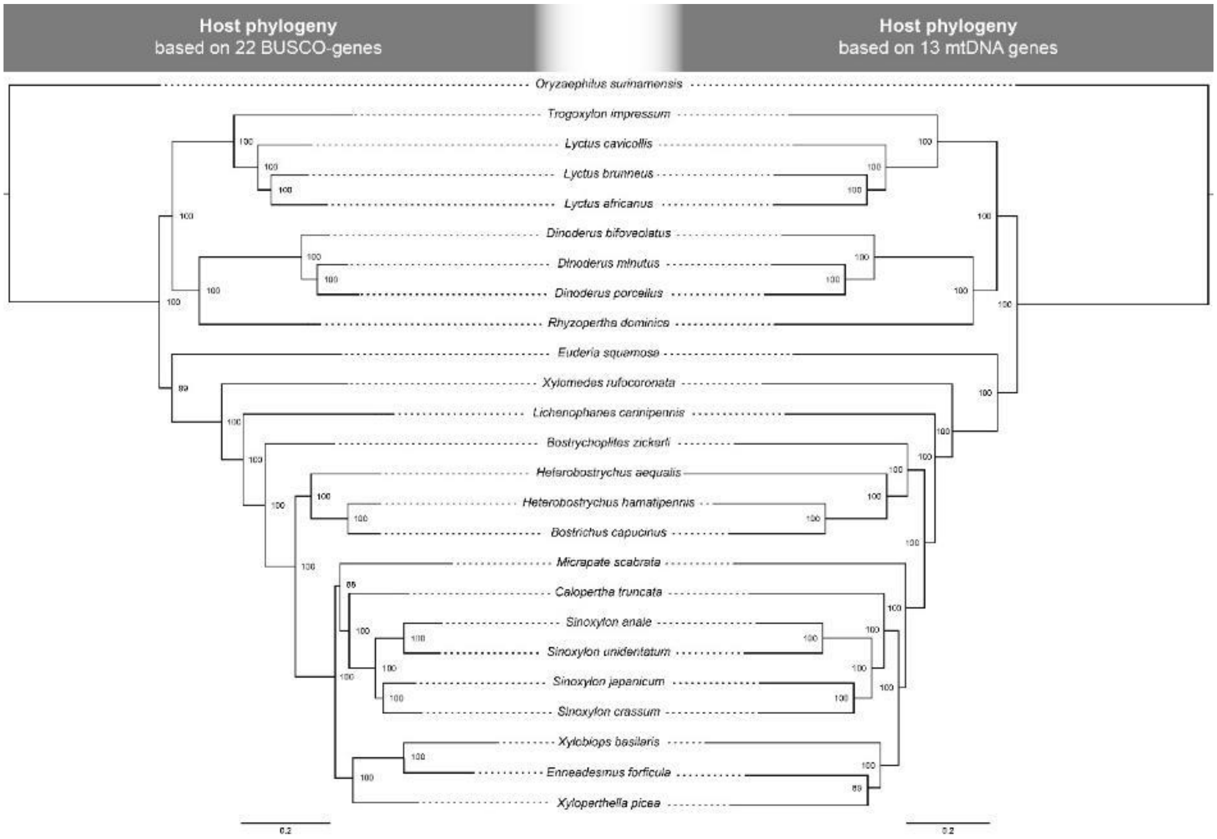


**Figure S2:** Comparison of host phylogenies based on 22 BUSCO genes (left) and 13 mitochondrial genes (right). Hosts with less than 22 annotated BUSCO genes were omitted from the analysis to achieve higher phylogenetic resolution. Node numbers represent posterior probabilities of Bayesian analyses.

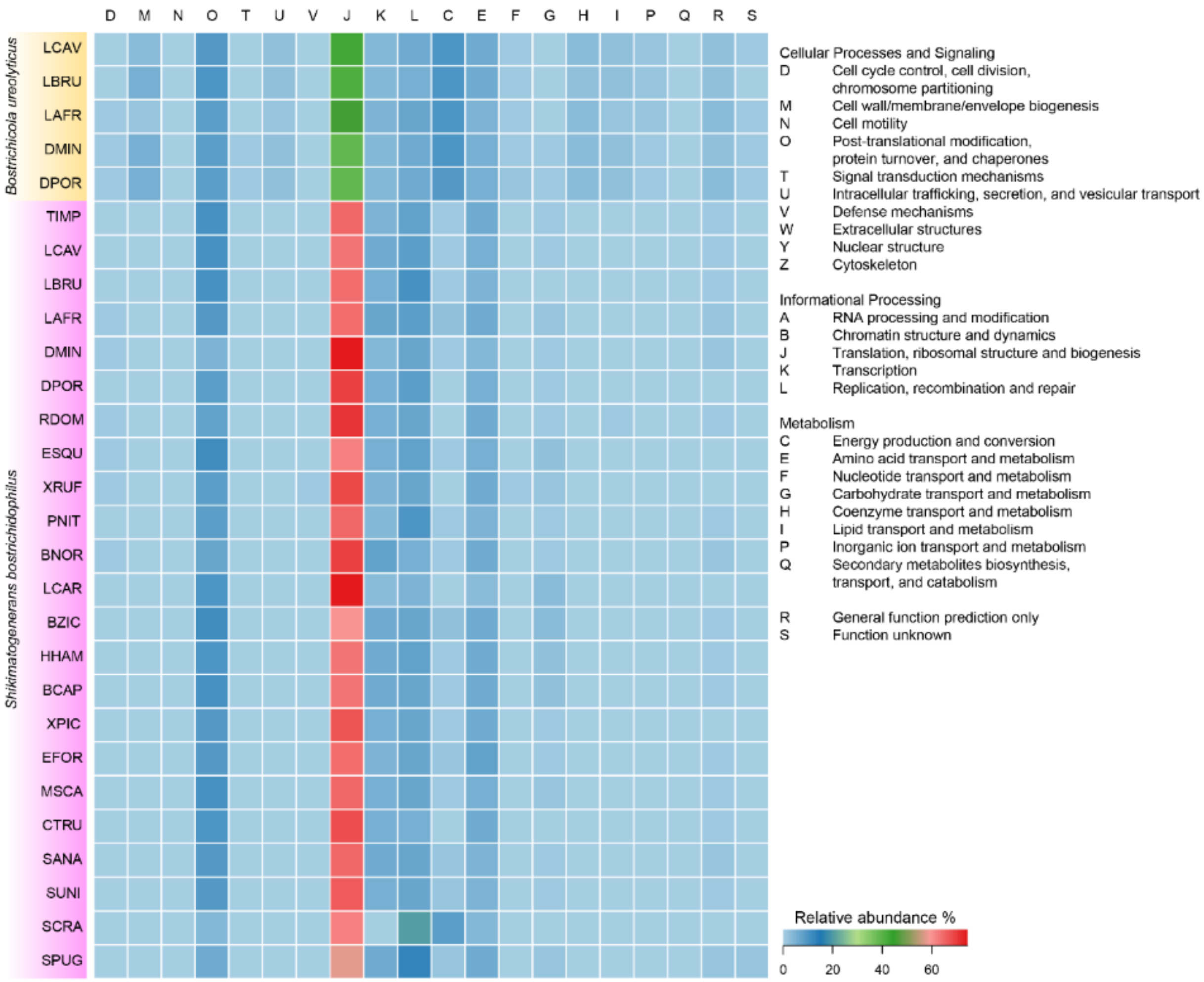


**Figure S3:** Relative abundance of Clusters of Orthologous Groups (COG). Annotated functional categories (A-Z) and relative proportion of the encoded genes represented as a heatmap are indicated on the right-hand side.

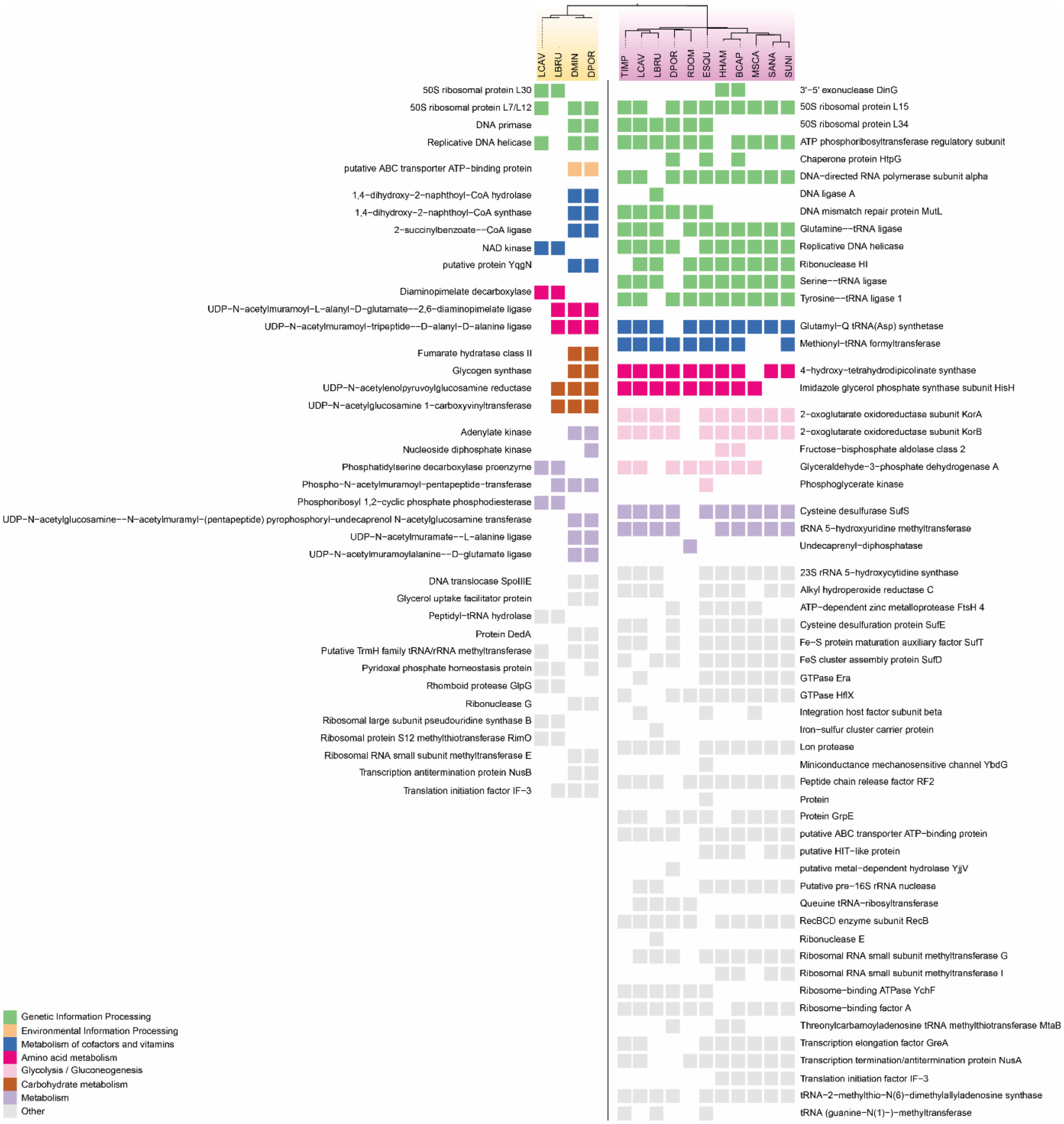


**Figure S4:** Comparison of all differentially encoded genes between genomes of *Bostrichicola ureolyticus* (left) and *Shikimatogenerans bostrichidophilus* (right). Genomes were annotated with PROKKA. Hypothetical proteins were removed. Filled: present, white: missing. Box colours are based on KEGG's categories (see legend for depicted categories)
